# Reduced emerin expression marks gastric cancer progression and invasive nuclear remodeling

**DOI:** 10.64898/2026.10.09.757948

**Authors:** Emily Hansen, Bovey Liu, James M. Holaska

**Author notes:** Correspondence: James M. Holaska, Cooper Medical School of Rowan University, Department of Biomedical Sciences, 401 South Broadway, Camden, NJ, 08103, USA.

## Abstract

Altered nuclear morphology is a defining diagnostic feature of malignancy, and nuclear structure is increasingly recognized as a functional regulator of cancer cell invasion. The inner nuclear membrane protein emerin maintains nuclear architecture and restrains invasive phenotypes in breast cancer models, but whether emerin expression is similarly altered in gastric cancer has not been established. Here, we analyzed emerin protein expression, nuclear morphology, and impeded migration in nonmalignant and malignant gastric cell lines. In parallel, emerin protein expression and nuclear envelope localization across gastric cancer was determined using patient tissue microarrays. Emerin was abundant at the nuclear envelope in cancer-adjacent and noncancerous gastric epithelium but was reduced across multiple gastric cancer histologies, including adenocarcinoma, signet-ring cell carcinoma, mucinous adenocarcinoma, neuroendocrine carcinoma, squamous cell carcinoma, and metastatic lesions. Reduced emerin at the nuclear envelope was most strongly associated with higher-grade tumors, with grade 2 and grade 3 cancers exhibiting significantly lower nuclear envelope scores than noncancerous tissue and grade 1 tumors. Consistent with the patient tissue data, the gastric cancer cell lines AGS and MKN1 expressed less emerin protein than noncancerous gastric epithelial control cells HFE-145. Consistent with studies in breast cancer cells, gastric cells with reduced emerin expression also had smaller nuclei and increased impeded migration through 8 µm pores. These findings support a model in which reduced nuclear envelope emerin is a conserved feature of invasive epithelial tumor progression and a candidate biomarker of gastric cancer nuclear remodeling.

## BACKGROUND

Gastric cancer remains a major cause of cancer morbidity and mortality worldwide and continues to present substantial therapeutic challenges because many patients are diagnosed with locally advanced or metastatic disease[1, 2]. Gastric tumors are also histologically and molecularly heterogeneous. Conventional pathological classifications distinguish intestinal, diffuse, signet-ring, mucinous, neuroendocrine, squamous, and mixed histologies, whereas molecular profiling has subdivided gastric adenocarcinoma into Epstein-Barr virus-positive, microsatellite unstable, genomically stable, and chromosomal instability-enriched groups[2, 3]. This heterogeneity complicates biomarker development and highlights the need for conserved cellular features that stratify tumor progression across histologic categories.

One of the most conserved features of malignant progression is altered nuclear morphology[4–7]. These structural changes are not merely diagnostic, as this altered morphology facilitates invasive behavior, which is considered one of the ‘hallmarks of cancer’[8, 9]. The nucleus is the largest and stiffest organelle (∼10-20 μm in diameter)[10], and its deformation is often rate limiting during metastasis because it requires confined migration through dense extracellular matrix and through small gaps in the vascular endothelium (∼1-2 μm in diameter)[11–14]. Thus, mechanisms that alter nuclear stiffness, nuclear envelope integrity, and nuclear-cytoskeletal coupling can influence whether tumor cells are capable of invasion, intravasation, extravasation, and metastatic colonization.

Emerin is a LEM-domain protein of the inner nuclear membrane that binds both nucleoskeletal components, such as nuclear lamins and actin, and chromatin regulatory proteins[15]. Through these interactions, emerin contributes to nuclear shape, nuclear mechanics, mechanotransduction, chromatin organization, and gene regulation[10, 15–29]. Prior work showed that invasive breast cancer cell lines have reduced emerin expression and smaller dysmorphic nuclei, resulting in their increased migration and invasion[30, 31]. Expression of wild type emerin in the invasive breast cancer cell lines rescued nuclear structure and reduced migration, invasion, tumor growth, and lung metastasis[31]. Expression of emerin mutants defective in nucleoskeletal binding failed to rescue these phenotypes, suggesting it was the nucleoskeletal interactions that were important. More recently, we showed that downregulating emerin in poorly invasive MCF7 breast cancer cells was sufficient to reduce nuclear size, increase nuclear deformation, and promote impeded migration[32]. In breast cancer patient tissue microarrays, emerin expression at the nuclear envelope inversely correlated with tumor invasiveness, stage, proliferation, and loss of differentiation markers[32, 33].

Emerin dysregulation is not restricted to breast cancer. Data from other cancers such as Ewing Sarcoma[34], melanoma,[35] and prostate cancer[36, 37] show that emerin protein reduction is associated with tumor transformation supporting a broader model in which reduced emerin expression or mislocalization weakens nuclear structural integrity, alters mechanotransduction, and facilitates invasive behavior[32, 33, 38]. However, the relevance of emerin to gastric cancer has not been defined. Using patient tissue microarrays, we show that emerin staining at the nuclear envelope is reduced across multiple gastric cancer histologies and in metastatic lesions. Using gastric epithelial and gastric cancer cell lines, we show that reduced emerin protein expression is associated with smaller, dysmorphic nuclei and increased impeded migration. These findings identify reduced emerin protein as a candidate marker of gastric cancer progression and support further mechanistic testing of emerin as a regulator of gastric cancer invasiveness.

## RESULTS

### Emerin expression at the nuclear envelope is reduced across gastric cancer histologic subtypes and metastatic lesions

To determine whether emerin expression is altered in gastric cancer patient tissue, we stained gastric cancer tissue microarrays containing 327 samples for emerin and scored nuclear envelope-localized staining using the same 0-3 nuclear envelope scoring strategy used in our breast cancer tissue microarray studies[32, 33]. A score of 0 corresponds to no detectable nuclear envelope rim staining, whereas a score of 3 corresponds to strong, continuous nuclear envelope staining. This scoring strategy measures both emerin abundance and appropriate nuclear envelope localization.

Cancer-adjacent/noncancerous gastric epithelium showed robust emerin staining at the nuclear envelope, with a mean nuclear envelope score of 2.07 ± 0.81 (Figure 1A). In contrast, all gastric cancers examined showed lower and more heterogeneous nuclear envelope staining of emerin. Squamous cell carcinoma (0.875 ± 0.89), neuroendocrine carcinoma (0.938 ± 1.3), and signet-ring carcinoma (0.958 ± 0.68) showed the lowest nuclear envelope scores. Adenocarcinoma (1.38 ± 0.95) and mucinous adenocarcinoma (1.28 ± 1.2) also showed significantly reduced emerin staining compared with cancer-adjacent/noncancerous tissue. Unsurprisingly, metastatic lesions showed significantly reduced emerin staining (0.982 ± 0.85) (Figure 1A). Representative images showed strong nuclear envelope staining in cancer-adjacent glands, whereas malignant tissues and metastatic lesions frequently showed weaker, discontinuous, or cytoplasmic/background staining patterns with less distinct nuclear rim enrichment (Figure 1B).

**Figure 1.**
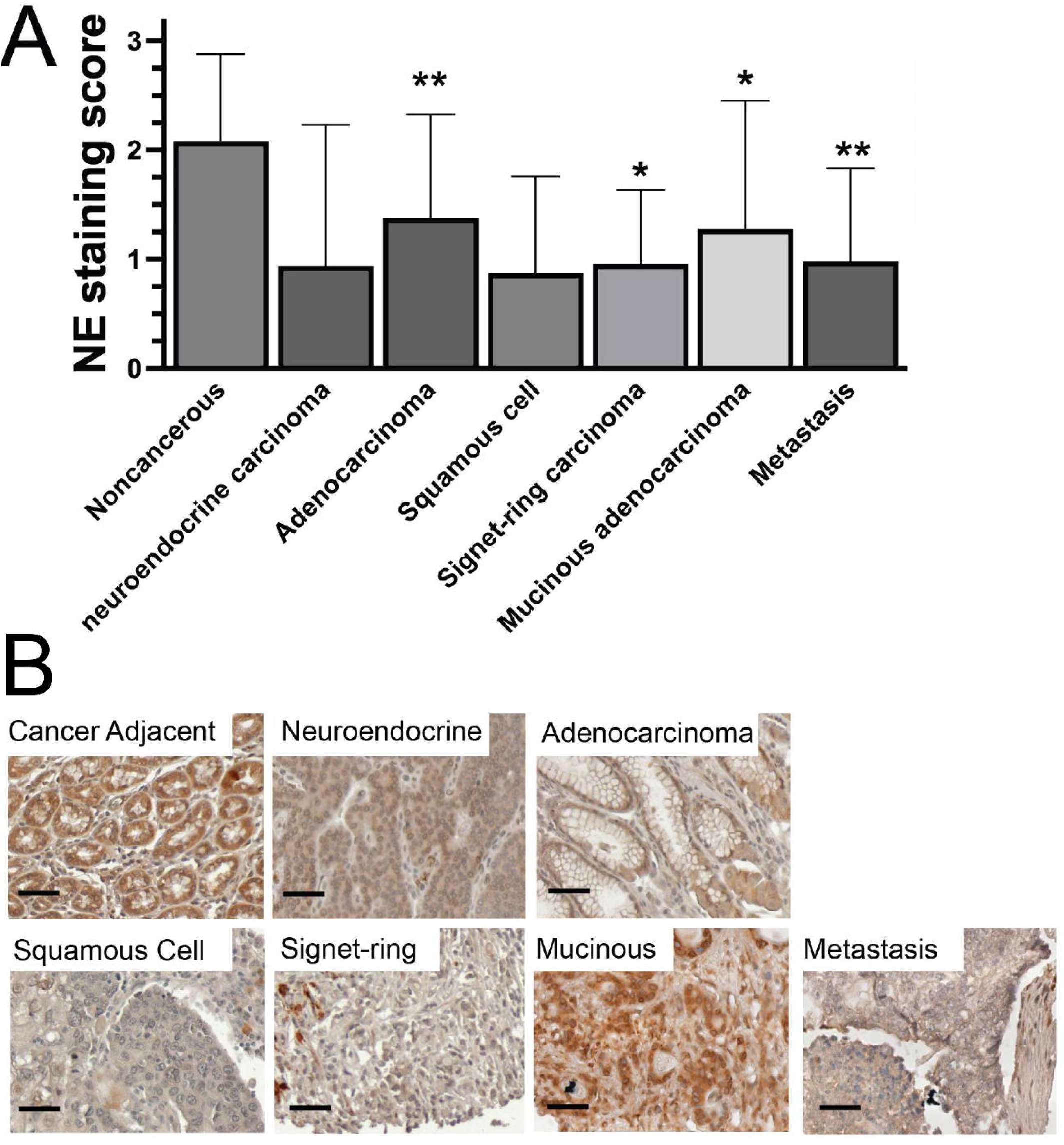
Reduced emerin expression at the nuclear envelope across gastric cancer histologic subtypes and metastatic lesions. A) Quantification of emerin nuclear envelope staining in 327 patient samples from cancer-adjacent/noncancerous gastric tissue, neuroendocrine carcinoma, adenocarcinoma, squamous cell carcinoma, signet-ring cell carcinoma, mucinous adenocarcinoma, and metastatic gastric cancer. Nuclear envelope staining was scored on a 0-3 scale, with 0 indicating no nuclear envelope staining and 3 indicating strong continuous nuclear rim staining. N=327, *P < 0.03, **P = 0.0005 compared to non-cancerous samples using one-way ANOVA followed by Dunnett’s multiple comparisons. B) Representative immunohistochemistry images showing emerin staining in the indicated samples using emerin polyclonal antibodies (Proteintech, cat# 10351-1-AP) or secondary alone (Vector Lab, cat#: MP-7451). Nuclei are blue and emerin is brown. As severity of cases increases, there is a visible reduction in emerin expression at the nuclear envelope and more deformed nuclei are present. Scale bars are 50 microns.

These data indicate that reduced emerin at the nuclear envelope is common across multiple gastric cancers rather than being restricted to a single subtype. These findings, including emerin reduction at the nuclear envelope in metastatic lesions, are consistent with the model that emerin loss marks, and may contribute to, invasive tumor progression.

### Emerin nuclear envelope staining is reduced in higher-grade gastric cancers

We next asked whether reduced emerin expression correlated with pathological grade or stage. 222 of the 327 samples were graded and used for this analysis, along with 69 noncancerous gastric tissue samples. Noncancerous gastric tissue (1.87 ± 0.90) and grade 1 gastric tumors (2.02 ± 0.74) retained relatively high emerin nuclear envelope staining (Figure 2A). In contrast, grade 2 (1.24 ± 0.88) and grade 3 (1.06 ± 0.88) cancers showed significantly lower emerin staining (Figure 2A). Representative tissue images are shown illustrating prominent nuclear rim staining in noncancerous and grade 1 samples, with weaker nuclear envelope staining of emerin and more irregular tumor nuclei in grade 2 and grade 3 specimens (Figure 2B).

**Figure 2.**
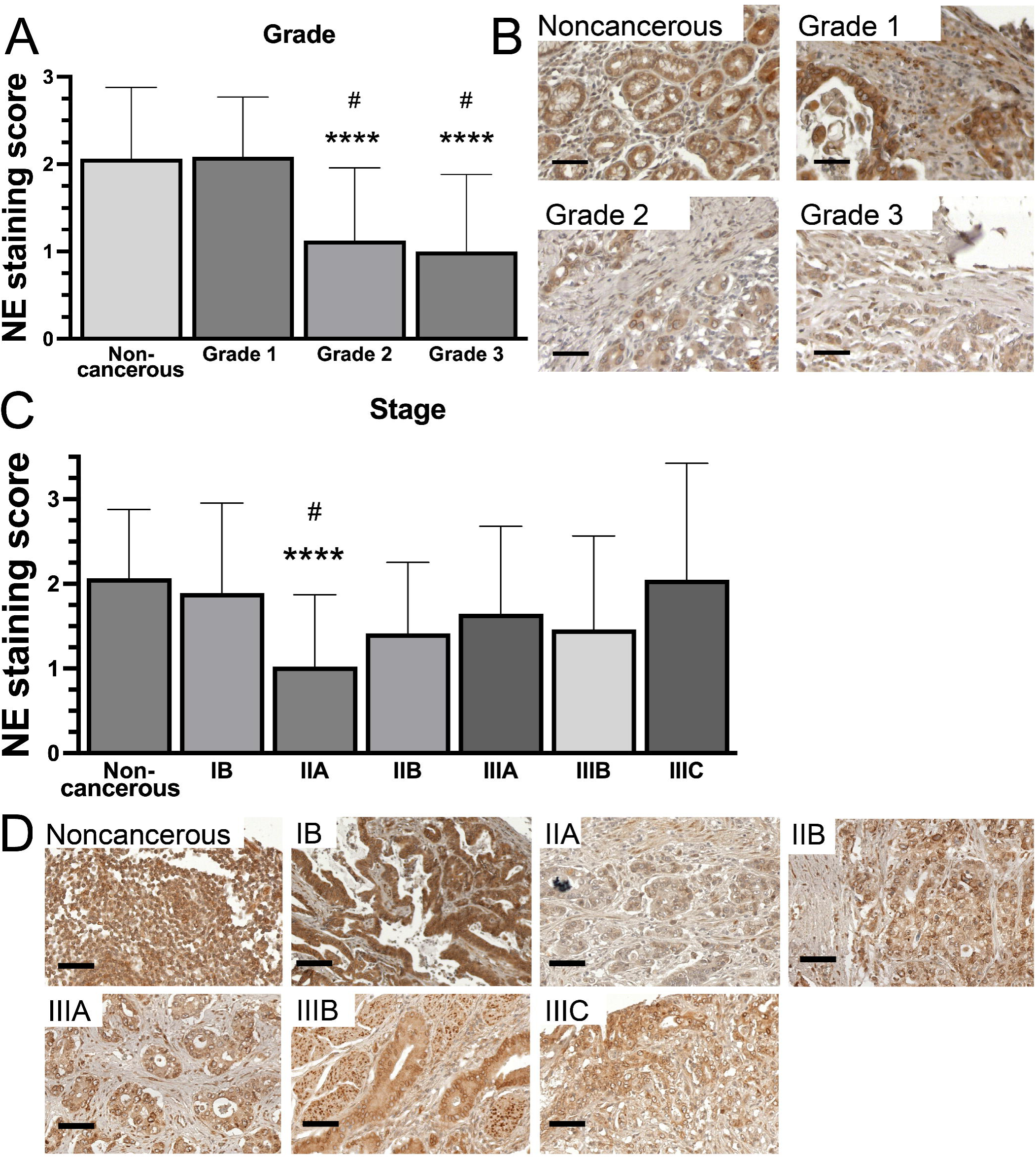
Emerin expression is reduced in higher-grade gastric cancers and is heterogeneous across stage. A) Quantification of the 222 tumor-graded tissue samples and 69 noncancerous gastric tissue samples for emerin nuclear envelope staining in which nuclear envelope staining was scored on a 0-3 scale, with 0 indicating no nuclear envelope staining and 3 indicating strong continuous nuclear rim staining . N=291 samples, ****P < 0.0001 compared to non-cancerous samples; ^#^P<0.002 compared to Grade 1 samples using one-way ANOVA followed by Tukey’s multiple comparisons. B) Representative emerin immunohistochemistry images for noncancerous, grade 1, grade 2, and grade 3 gastric tissue using emerin polyclonal antibodies (Proteintech, cat# 10351-1-AP) or secondary alone (Vector Lab, cat#: MP-7451). Nuclei are blue and emerin is brown. As tumor grade increases, there is a visible reduction in emerin expression at the nuclear envelope and more deformed nuclei are present. Scale bars are 50 microns. C) Quantification of emerin nuclear envelope staining in the 125 tumor-staged and 69 noncancerous tissue samples. N=194 samples, ****P < 0.0001 compared to non-cancerous samples, ^#^P=0.0179 compared to Stage IB samples using one-way ANOVA followed by Tukey’s multiple comparisons. D) Representative emerin immunohistochemistry images for noncancerous tissue and indicated stages using emerin polyclonal antibodies (Proteintech, cat# 10351-1-AP) or secondary alone (Vector Lab, cat#: MP-7451). Scale bars are 50 microns.

Of the 327 samples, 125 samples had available staging information and were analyzed for association of reduced emerin protein with cancer staging as compared to 69 noncancerous gastric tissue samples. When samples were stratified by stage, emerin staining was more heterogeneous. Stage IB gastric tumors showed emerin nuclear envelope scores (2.09 ± 0.89) comparable to noncancerous tissue (2.07 ± 0.81), whereas stage IIA tumors showed a marked reduction in nuclear envelope emerin (1.06 ± 0.82) (Figure 2C). Stage IIB (1.46 ± 1.1), IIIA (1.62 ± 0.98), and IIIB (1.51 ± 0.90) tumors trended lower, but more samples will be needed to determine significance for these stages, for which there were 2-10-fold less samples compared to stage IIA tumors represented on the tissue microarrays (Table S1). Although stage IIIC samples had higher nuclear envelope staining scores for emerin (2.05 ± 1.38), there was substantial variability due to low sample numbers (n=5). Representative stage-matched images (Figure 2D) show the differences in both staining intensity of emerin at the nuclear envelope and nuclear morphology across stages (Figure 2D). Collectively, these data suggest that loss of emerin at the nuclear envelope is associated most strongly with grade, tumor progression, and metastasis in this gastric cancer cohort.

### Gastric cancer cell lines express less emerin protein than noncancerous gastric epithelial controls

To test whether the reduction in emerin observed in gastric cancer tissues was mirrored in gastric cancer cell lines, we analyzed emerin protein expression in the noncancerous gastric epithelial cell line HFE-145 and in the gastric cancer cell lines AGS and MKN1 by western blotting (Figure 3A). Emerin levels were normalized to β-actin and shown relative to HFE-145 cells (Figure 3B). HFE-145 cells showed the highest emerin protein expression and were set to 1.0 (1.00 ± 0.12, n=3). Both AGS (0.292 ± 0.13, n=3) and MKN1 (0.413 ± 0.18, n=3) gastric cancer cell lines had significantly reduced emerin expression (Figure 3B). These data show that reduced emerin protein expression is a reproducible feature of invasive gastric cancer cell lines, consistent with reduced nuclear envelope emerin staining observed in patient specimens.

**Figure 3.**
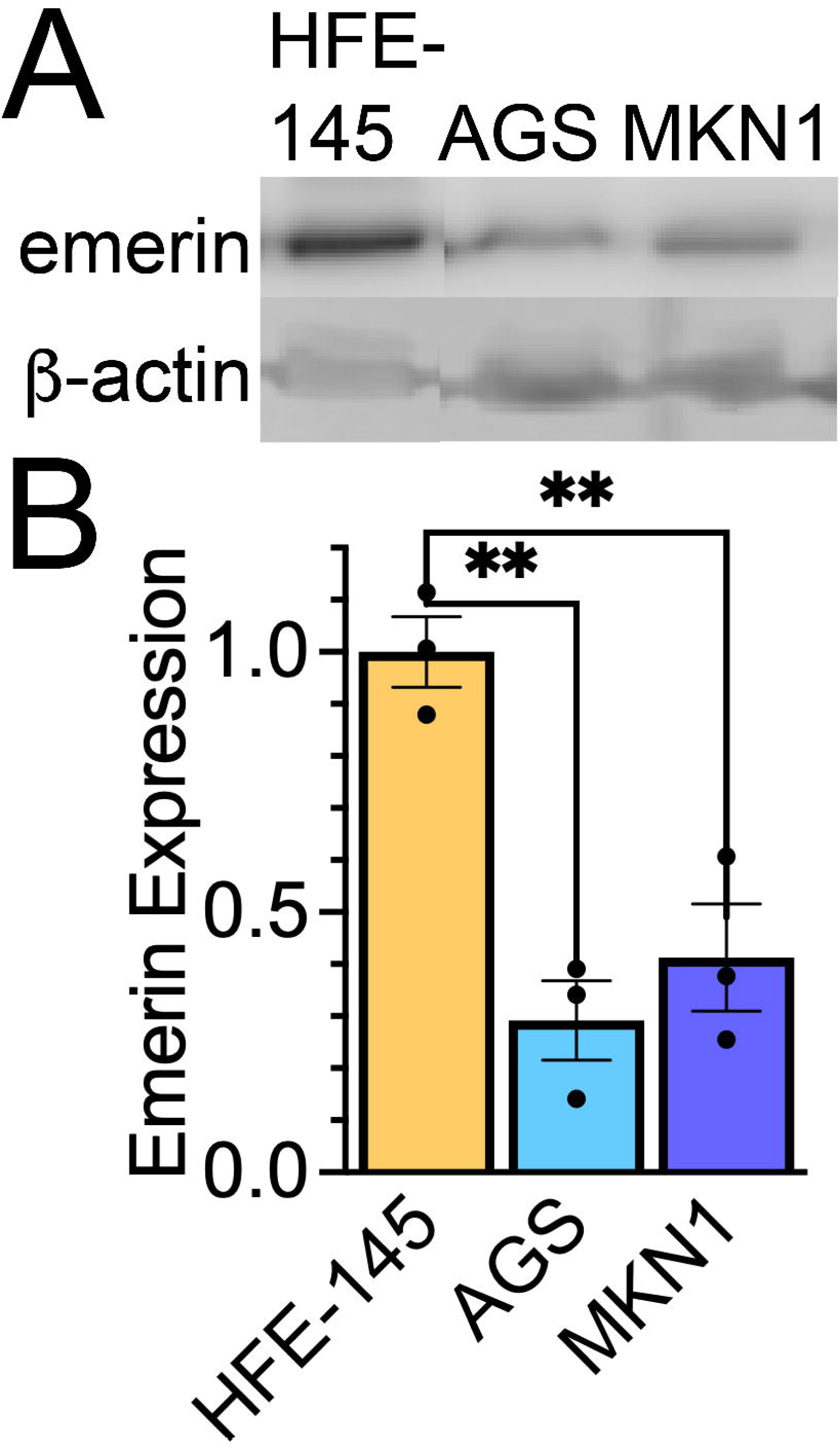
Gastric cancer cell lines show reduced emerin protein expression compared with noncancerous gastric epithelial cells. A) Representative western blot of emerin and β−actin in HFE-145, AGS, and MKN1 cells. B) Quantification of emerin protein expression normalized to β−actin and expressed relative to HFE-145 cells. Dots represent independent biological replicates. Error bars represent SD, n=3. **P<0.005, One-way ANOVA followed by Dunnett’s multiple comparisons.

### Invasive gastric cancer cell lines have smaller nuclei and increased impeded migration

Emerin maintains nuclear structure in multiple cell types, and reduced emerin expression in breast cancer models decreases nuclear size and increases nuclear dysmorphism. We therefore asked whether AGS and MKN1 gastric cancer cell lines, which express less emerin, also exhibited altered nuclear morphology. DAPI-stained nuclei were imaged, and nuclear area was quantified using the Image J ParticleAnalyzer plug-in. HFE-145 cells had larger nuclei (213.2 ± 73.8 µm^2^, n=116) than AGS (80.32 ± 29.9 µm^2^, n=249) and MKN1 (80.43 ± 27.7 µm^2^, n=183) (Figure 4A,B). These data show that reduced emerin protein expression is associated with smaller nuclei in gastric cancer cells, consistent with previous findings in breast cancer and prostate cancer models[31, 32, 36, 37, 39, 40].

**Figure 4.**
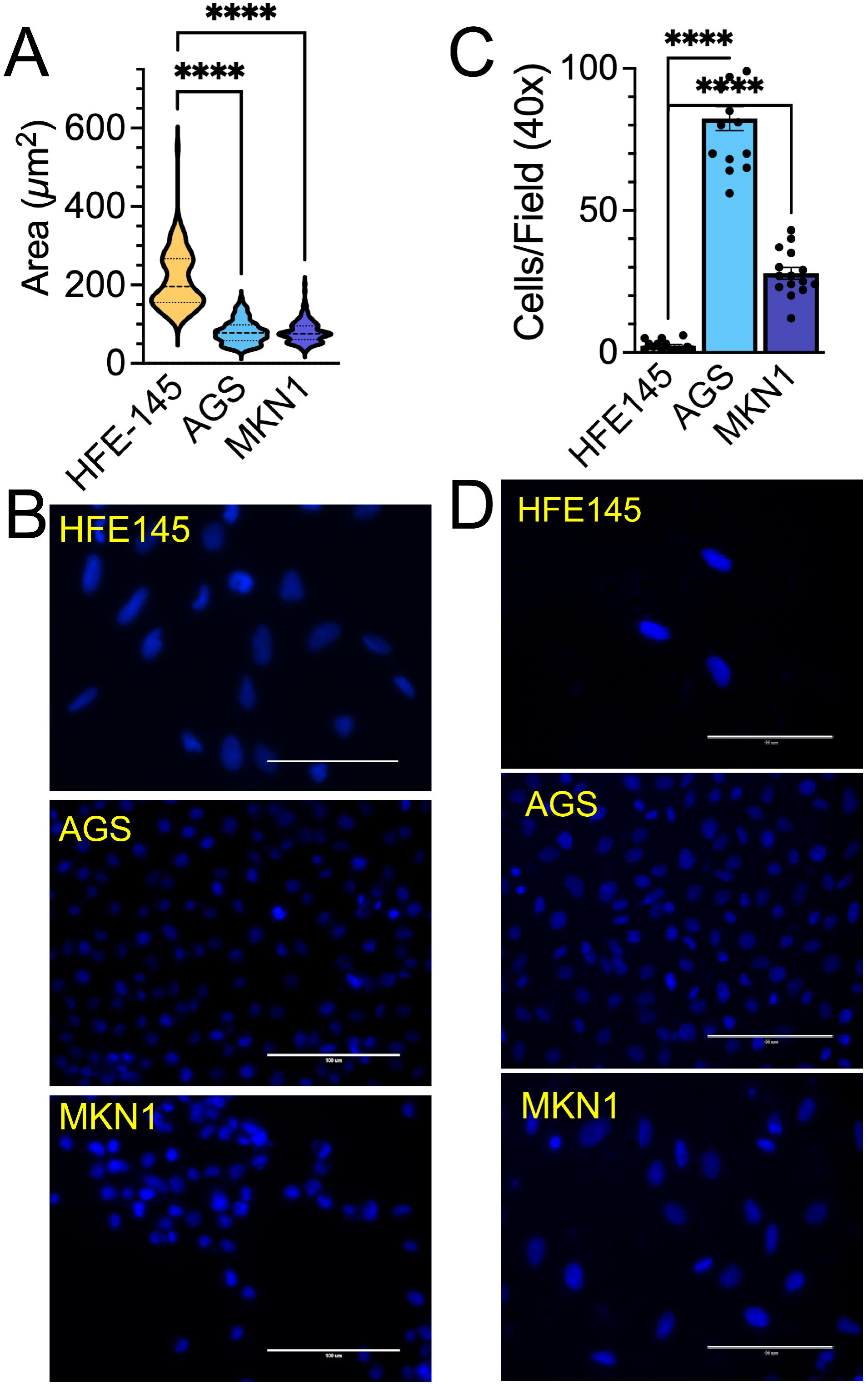
Gastric cancer cell lines with reduced emerin expression have smaller nuclei and increased impeded migration. A) Violin plot of nuclear area of HFE-145, AGS, and MKN1 cells (N>115 nuclei). The mean is depicted as the dark dashed line and the thin dashed lines represent the first and third quartiles. ****P<0.0001, one-way ANOVA followed by Dunnett’s multiple comparisons. B) Representative DAPI images of HFE-145, AGS, and MKN1 cells, which were used to measure nuclear area. Scale bars are 100 microns. C) Quantification of cells migrating through 8 micrometer trans-well pores, expressed as cells per field at 40x magnification (n=15 fields). Dots represent independent biological replicates. Error bars represent SD. ****P<0.0001, one-way ANOVA followed by Tukey’s multiple comparisons. D) Representative DAPI images of cells that successfully migrated. Scale bars are 100 microns.

Thus, we tested whether AGS or MKN1 cells exhibited increased impeded migration through 8 µm transwell pores. AGS cells showed the highest impeded migration, with 82.33 ± 16.4 cells per 40x field (n=15), while in MKN1 cells, 27.87 ± 8.14 cells per 40x field (n=15) migrated through the transwell (Figure 4 C,D). HFE-145 and MKN45 cells migrated poorly, with only 2.400 ± 1.72 cells per 40x field (n=15) migrated through 8 µm transwell pores. Thus, reduced emerin protein expression correlated with increased impeded migration in AGS and MKN1 cells.

## DISCUSSION

This study identifies reduced emerin protein expression at the nuclear envelope as a feature of gastric cancer progression. Across gastric cancer tissue microarrays, emerin staining at the nuclear envelope was lower in multiple malignant samples than in cancer-adjacent/noncancerous gastric epithelium. Reduced emerin staining was also associated with higher tumor grade. In parallel, gastric cancer cell lines expressed less emerin protein and had reduced emerin expression, smaller nuclei, and increased impeded migration compared to noncancerous gastric epithelial controls. These results extend our previous breast cancer findings and suggest that emerin loss may represent a conserved nuclear envelope alteration during epithelial cancer progression.

In patient tissues, the strongest association was between reduced emerin protein and tumor grade. Grade 2 and grade 3 tumors exhibited lower emerin expression at the nuclear envelope compared to grade 1 tumors. This pattern is consistent with a model in which emerin loss accompanies tumor dedifferentiation and/or preferential selection for cells with altered nuclear architecture to facilitate invasion and migration. The stage analysis suggests that emerin reduction may occur in subsets of clinically advanced disease, but larger stage-balanced cohorts will be needed to determine whether emerin independently stratifies with stage, lymph node involvement, metastatic dissemination, or survival. In breast cancer, emerin reduction correlated with increased grade and molecular features of aggressive disease, including advanced stage, increased Ki-67 expression, loss of ER/PR expression, and receptor-negative phenotypes[32, 33]. Though gastric cancer is not routinely stratified using a receptor-based clinical system, histologic grade and subtype capture aspects of differentiation, invasiveness, and tissue organization. The finding that reduced emerin spans diverse gastric cancer tumors, including adenocarcinoma, signet-ring, mucinous, neuroendocrine, squamous, and metastatic samples suggests that emerin reduction may represent a shared feature important for driving gastric tumorigenesis.

A key implication of these findings is that nuclear envelope remodeling may contribute to gastric cancer invasion. The nucleus poses a physical barrier during migration through confined spaces, and tumor cells must deform their nuclei to invade dense extracellular matrix, transit through small gaps in the vascular endothelium, and to colonize distant tissues[11–14]. Emerin interacts with lamins and nuclear actin and contributes to nuclear stiffness, nuclear shape, chromatin organization, and mechanotransduction[7, 15]. In breast cancer models, loss of emerin reduced nuclear size, increased nuclear deformation, and promoted migration through small pores, whereas re-expression of emerin restored nuclear structure and suppressed invasion and metastasis[30–32, 38]. The gastric cancer data presented here are consistent with this model: gastric cancer lines with reduced emerin generally had smaller nuclei and exhibited increased migration through 8 µm pores. These results support a model in which reduced emerin promotes nuclear dysmorphism to facilitate migration of invasive cancer cells through confined spaces.

These findings have several potential translational implications. First, emerin nuclear envelope staining could be developed as a pathology-compatible biomarker of nuclear remodeling in gastric cancer. Tissue microarray immunohistochemistry can be integrated with routine histology, and the nuclear rim staining pattern is interpretable in formalin-fixed paraffin-embedded samples. Second, reduced emerin may help identify tumors with altered nuclear mechanics that are more capable of confined migration, metastatic dissemination, or resistance to mechanical stress. Third, emerin-associated pathways may represent therapeutic vulnerabilities in invasive gastric cancer. Combined with previous work showing emerin’s importance in breast and prostate cancer, this work supports targeting emerin-associated pathways as therapeutic targets.

The present study also has limitations. The tissue microarray analysis is semi-quantitative and correlative. Further, larger independent cohorts with more comprehensive patient data will be required to evaluate associations of emerin reduction with lymph node status, metastatic site, recurrence, survival, and therapy response. Future studies beyond the scope of this work, will need to determine the effects of emerin knockdown or knockout in noninvasive gastric epithelial cells, emerin restoration in emerin-low gastric cancer cells, and test emerin mutants defective in nucleoskeletal binding, as well as binding to chromatin-associated proteins and mechanotransduction machinery.

## CONCLUSION

In conclusion, reduced emerin expression at the nuclear envelope is associated with increased gastric cancer aggressiveness and histologic grade, reduced nuclear size, and increased impeded migration. In gastric cancer cell lines, lower emerin expression was also associated with reduced nuclear area and increased impeded migration. These results support a model in which emerin loss contributes to a conserved nuclear envelope remodeling program that promotes invasive potential in epithelial cancers. Further mechanistic studies will determine whether emerin is simply a biomarker of gastric cancer progression or a targetable suppressor of metastatic transformation.

## METHODS

### Immunohistochemistry and Tissue Microarray Analysis

All patient samples used for these studies are provided as tissue microarrays and were not collected by our group; rather they were obtained from a commercial source (TissueArray, LLC) that provides IHC-ready histological samples directly on slides. Age, grading, staging, pathology diagnosis, type, and molecular phenotyping were provided with each sample (Table S1). Tissue microarrays (TissueArray, LLC, cat# ST810d, ST1451a, and T011B) for emerin-stained samples and cat# ST8013b for secondary-only tissue) were deparaffinized and rehydrated in Coplin jars (4 minutes in xylene three times, 3 minutes in 100% EtOH two times, 3 minutes in 95% EtOH two times, 3 minutes in 80% EtOH, 3 minutes in 70% EtOH, and 5 minutes in distilled water) prior to being placed in citrate buffer (0.05% Tween 20, 10mM citric acid at pH 6.0) and steamed at 95°C for 45 minutes. Slides were cooled at RT and washed with PBS. Endogenous peroxidase was removed by washing with 0.3% hydrogen peroxide for 20 minutes at room temperature in coplin jars. After washing again with PBS, slides were blocked in 1% Bovine Serum Albumin (VWR, cat#: 97061-420) in PBS for 15 minutes. Slides were then incubated with anti-emerin antibody (Proteintech, cat#: 10351-1-AP, 1:500 dilution) for two hours at 37°C in a humidified chamber. Slides were washed again with PBS and blocked with 2.5% Normal Goat Serum for 20 minutes at room temperature (ImmPRESS Reagent, Vector Lab, cat#: MP-7451) and then incubated with the anti-rabbit ImmPRESS IgG peroxidase reagent (Vector Lab, cat#: MP-7451) or the anti-mouse ImmPRESS IgG peroxidase reagent (Vector Lab, cat# MP-7452) per manufacturer instructions. Slides were washed again with PBS and incubated with the ImmPACT DAB peroxidase substrate (Vector Lab, cat#: SK4105) for 90 seconds at room temperature while checking color development under a microscope before rinsing in tap water. Slides were counterstained with Vector Hematoxylin (Gill’s Formula, Vector Lab, cat#: H3401) for three minutes. After rinsing again in tap water, slides were incubated in 0.1% sodium bicarbonate for one minute, rinsed with tap followed by distilled water, and then dehydrated and mounted with Prolong Diamond Antifade Mountant (Invitrogen, cat#: P36970). Tissue images were taken using the Evos FL Auto microscope and the Precipoint slide scanning microscope.

### Nuclear Envelope Score

The presence of all patient samples on a single slide serves as an internal control for antibody staining and the ability to accurately score samples relative to the control samples (e.g., noncancerous). This is important, as IHC staining protocols generally are optimized to ensure maximal staining and thus incubation times with substrates to produce the stain may vary between slides. A score of 3 was given for samples that had maximal staining of emerin localized exclusively at the nuclear envelope; note that noncancerous samples exhibited variability and did not always score a 3, as this was reserved for the highest levels of staining. A score of 2 represented reduced emerin staining with this staining at the nuclear envelope. A score of 1 represented reduced emerin staining that was not only at the nuclear envelope or emerin staining that was less than samples with a score of 2 that localized exclusively at the nuclear envelope. A score of 0 was given to those samples that lacked detectable emerin. The graders were blinded to sample identifiers; there were at least two graders for each. The presence of all samples on a single slide that were treated similarly, combined with the comparison of emerin protein levels to controls, allows us to measure protein expression in each sample relative to control. Further, the presence of non-cancerous controls on tissue microarrays ST1451A and T011b allowed for scoring of each patient tissue on a given tissue microarray to the control samples on that tissue microarray. Thus, comparisons between different tissue microarrays could be done because they were each internally controlled and compared to controls samples set at maximal score of 3, as described above. All grading was done in one sitting to avoid multi-day bias. Invasive, DCIS, and metastatic tissue measurements were determined significant against noncancerous tissue using one-way ANOVA with Tukey or Dunnett’s comparison, with the data presented as Mean ± SD.

### Cell Culture

HFE-145, AGS (ATCC cat#: ctrl-1739), and MKN1 cells were obtained from Shumei Song’s lab (Coriell Institute for Medical Research, Camden NJ). All cells were grown in Dulbecco’s Modified Eagle Medium + GlutaMAX (Gibco cat#:10566024) with 7% Fetal Bovine Serum (Gibco, cat#: 16140089) and 1% Penicillin/Streptomycin. All cells were grown at 37°C and 5% CO_2_. Mycoplasma testing was done bimonthly using the MycoStrip 100 kit (InvivoGen, cat#: rep-mysnc-100).

### Immunofluorescence, volume, and nuclear area measurement

Cell lines were plated on coverslips at approximately 100,000 cells/coverslip and placed in 6-well plates. Cells were rinsed three times with 2 ml of PBS for 5 minutes and fixed with 3.7% formaldehyde in PBS for 15 minutes. Coverslips were washed again three times and permeabilized in 2 ml of 0.2% Triton X-100 in PBS for 24 minutes. Cells were blocked for 1 hour in 2 ml of 3% BSA in PBS (VWR, cat#: 97063-624). Cells were then washed three times with 2 ml of PBS for 5 minutes each and mounted with Prolong Diamond Antifade Mountant with DAPI (Molecular Probes, cat#: P36971). Slides were imaged using either the Evos FL Auto Microscope using a 40x objective or Nikon confocal with 100x objective. Nuclear area was measured via Image J ParticleAnalyzer plugin. Briefly, images were transformed into a binary image where ParticleAnalyzer could then measure each nucleus. Nuclei on the edges of the image or touching other nuclei were excluded. Nuclear volume was measured via ImageJ FIJI 3D Objects Counter Plugin following the same parameters. Statistical significance was determined via one-way ANOVA followed by Dunnett’s multiple comparison test, where applicable.

### Transwell Migration Assays

Transwell inserts with 8-micron pores (Falcon, Cat#: 353097) were used for cell migration. Cells were plated in the chamber at a density of 1.5 x10^5^ in serum-free media. Chambers were placed in 24-well plates containing complete growth media. After 24 hours, cells that failed to migrate through the trans-well were wiped off the top of the membrane and the cells on the bottom of the membrane were fixed for 15 minutes with 3.7% formaldehyde in PBS. The membranes were then washed 3 times in 750 µl of PBS for 5 minutes and treated with 750 µl of 0.5% Triton X-100 for 20 minutes to permeabilize membranes. The membranes were then washed a final three times with PBS and removed and mounted, cell-side down, with Prolong Diamond Antifade Mountant with DAPI (Molecular Probes, cat#: P36971). Cells that migrated through were counted (five fields on the Evos FL Auto Microscope, 40x objective) after allowing the mountant to dry overnight. Statistical significance was determined via one-way ANOVA followed by Dunnett’s multiple comparison test, where applicable. At least three biological replicates were done for each cell type.

### Western Blots

Whole cell lysates were suspended in NuPAGE LDS Buffer (LifeTechnologies, cat#: NP0008) with reducing agent (LifeTechnologies, cat#: NP0009) at a concentration of 5×10^6^ cells/ml and heated 10 minutes at 70°C followed by brief sonication. 50,000 cells from each sample were resolved by 10% SDS-PAGE gels (materials from Bio-Rad) and transferred to nitrocellulose membranes (GE Healthcare, Cat#: 10600004). Membranes were blocked in 5% nonfat dry milk (Giant brand) in PBST for two hours. Primary emerin (Leica Biosystems; #NCL-EMERIN, 1:3,000) and β-actin (Sigma, cat# SAB5600204, 1:1,000) antibodies were incubated overnight with rocking at 4°C and secondary antibodies (goat anti-rabbit [cat# 31462, Invitrogen, 1:10,000 ] or goat anti-mouse [cat#31432, Invitrogen, 1:10,000] IgG H&L cross absorbed HRP) were incubated for 2 hours at room temperature. Emerin protein expression was normalized to β-actin protein expression for quantification. Western blots were imaged on the Li-COR Odyssey FC. Statistical significance was determined via one-way ANOVA.

### Statistical analysis

Statistical analyses were performed using GraphPad Prism. Data are presented as mean +/-SD unless otherwise noted. Comparisons among more than two groups were performed using one-way ANOVA followed by Dunnett or Tukey multiple-comparison tests, as appropriate. For tissue microarray analyses, cancer subtype, grade, and stage groups were compared with noncancerous or indicated reference groups and significance was determined using one-way ANOVA followed by Dunnett or Tukey multiple-comparison tests, as appropriate. Significance symbols in the figures should be defined as follows after confirmation: *P < 0.05, **P < 0.01, ***P < 0.001, ****P < 0.0001, ^#^P=0.01 compared to Grade 1 (Figure 2A) or Stage IB (Figure 2B), and ^###^P<0.0001.

## DATA AVAILABILITY

The data supporting the findings of this study are available from the corresponding author upon reasonable request.

## ETHICS APPROVAL AND CONSENT TO PARTICIPATE

All human tissue specimens were obtained as commercially available, de-identified tissue microarrays. The vendor states that tissues were collected under applicable ethical standards and consent procedures. No identifiable clinical data were collected by the authors. Please replace this statement with the exact vendor ethics language and institutional determination for the gastric tissue microarray used in this study.

## ACKNOWLEDGEMENTS

We thank the Department of Biomedical Sciences at Cooper Medical School of Rowan University for support and the members of the Holaska laboratory for many helpful discussions. We thank Dr. Shumei Song for sharing the gastric cancer cell lines used in this study.

## AUTHOR CONTRIBUTIONS

**Emily Hansen:** conceptualization; data curation; formal analysis; funding acquisition; investigation; methodology; validation; visualization; writing; reviewing and editing. **Bovey Liu:** data curation; formal analysis; investigation; validation. **James M: Holaska:** conceptualization; data curation; formal analysis; funding acquisition; investigation; methodology; project administration; resources; supervision; validation; visualization; writing; reviewing and editing.

## FUNDING

This work was supported by a grant from the National Institute of Arthritis, and Musculoskeletal and Skin Diseases (R15AR069935 to JH), a grant from the New Jersey Commission on Cancer Research (COCR22RBG007 to JH), and a grant from the New Jersey Commission on Cancer Research (COCR25PRF018 to EH). The content is solely the responsibility of the authors and does not necessarily represent the official views of the National Institutes of Health or the New Jersey Commission on Cancer Research. This work was also supported by the Camden Cancer Research Center and by Rowan University under the Camden Health Research Initiative.

## COMPETING INTERESTS

The authors declare no competing interests.

## Conflicts of Interest

Authors have no conflicts of interest to disclose.

## Data Availability

The data supporting the findings of this study are available by request from the corresponding author (J.M.H.).

## Ethics Approval and Consent to Participate

Not applicable

## Notes

### Competing Interest Statement

The authors have declared no competing interest.

