## Supplemental Table 1 for "Reduced emerin expression marks gastric cancer progression and invasive nuclear remodeling"

| Microarray | No. | Age | Sex | Organ/Anatomic Site | Pathology diagnosis | Grade | Stage | Type | Score |
|---|---|---|---|---|---|---|---|---|---|
| **ST810D** | | | | | | | | | |
| | A1 | 70 | M | Stomach | Adenocarcinoma | 1--2 | IIA | Malignant | 2.5 |
| | A2 | 71 | M | Stomach | Adenocarcinoma | 2 | IIA | Malignant | 1 |
| | A3 | 61 | M | Stomach | Adenocarcinoma | 2 | IIB | Malignant | 1 |
| | A4 | 60 | M | Stomach | Adenocarcinoma | 2 | IIA | Malignant | 0.5 |
| | A5 | 54 | M | Stomach | Adenocarcinoma | 3 | IIA | Malignant | 1.5 |
| | A6 | 72 | M | Stomach | Mucinous adenocarcinoma | 2 | IIA | Malignant | 0.5 |
| | A7 | 77 | M | Stomach | Adenocarcinoma | 2 | IIB | Malignant | 1 |
| | A8 | 49 | F | Stomach | Adenocarcinoma | 2 | IIB | Malignant | 1.5 |
| | A9 | 63 | M | Stomach | Adenocarcinoma | 2 | IIA | Malignant | 0.5 |
| | A10 | 66 | M | Stomach | Adenocarcinoma | 2 | IIA | Malignant | 0.25 |
| | A11 | 53 | M | Stomach | Adenocarcinoma | 2 | IIB | Malignant | 0 |
| | A12 | 68 | F | Stomach | Adenocarcinoma | 2 | IIA | Malignant | 1.25 |
| | A13 | 64 | F | Stomach | Adenocarcinoma | 2 | IIA | Malignant | 0.5 |
| | A14 | 56 | M | Stomach | Adenocarcinoma | 2 | IIB | Malignant | 2.5 |
| | A15 | 57 | M | Stomach | Adenocarcinoma | 2--3 | IIB | Malignant | 3 |
| | A16 | 47 | F | Stomach | Adenocarcinoma | 2--3 | IIB | Malignant | 0.5 |
| | A17 | 68 | M | Stomach | Adenocarcinoma | 2--3 | IIA | Malignant | 1.5 |
| | A18 | 56 | M | Stomach | Adenocarcinoma | 3 | IIA | Malignant | 3 |
| | A19 | 62 | M | Stomach | Adenocarcinoma | 3 | IIB | Malignant | 3 |
| | A20 | 68 | F | Stomach | Adenocarcinoma | 3 | IIA | Malignant | 1.5 |
| | A21 | 36 | M | Stomach | Mucinous adenocarcinoma | 3 | IIB | Malignant | 0.25 |
| | A22 | 53 | M | Stomach | Adenocarcinoma | 3 | IIA | Malignant | 0 |
| | A23 | 54 | M | Stomach | Adenocarcinoma | 3 | IIB | Malignant | 0 |
| | A24 | 34 | M | Stomach | Adenocarcinoma | 3 | IIA | Malignant | 0.5 |
| | A25 | 45 | M | Stomach | Adenocarcinoma | 3 | IIA | Malignant | 1.5 |
| | A26 | 49 | M | Stomach | Adenocarcinoma | 3 | IIA | Malignant | 1 |
| | A27 | 57 | M | Stomach | Adenocarcinoma | 3 | IIB | Malignant | 3 |
| | A28 | 63 | M | Stomach | Adenocarcinoma | 3 | IIIA | Malignant | 3 |
| | A29 | 59 | M | Stomach | Adenocarcinoma | 3 | IIA | Malignant | 3 |
| | A30 | 60 | M | Stomach | Adenocarcinoma | 3 | IIA | Malignant | 2.5 |
| | A31 | 47 | F | Stomach | Adenocarcinoma | 3 | IIB | Malignant | 2 |
| | A32 | 50 | M | Stomach | Adenocarcinoma | 3 | IIA | Malignant | 1 |
| | A33 | 63 | M | Stomach | Adenocarcinoma | 3 | IIA | Malignant | 3 |
| | A34 | 37 | F | Stomach | Adenocarcinoma | 3 | IIB | Malignant | 1 |
| | A35 | 35 | M | Stomach | Adenocarcinoma | 3 | IIB | Malignant | 1 |
| | A36 | 68 | M | Stomach | Mucinous adenocarcinoma | 3 | IIA | Malignant | 2 |
| | A37 | 58 | M | Stomach | Adenocarcinoma | 3 | IIA | Malignant | 2 |
| | A38 | 24 | F | Stomach | Adenocarcinoma | 3 | IIA | Malignant | 2.5 |
| | A39 | 64 | M | Stomach | Adenocarcinoma | 3 | IIA | Malignant | 2.5 |
| | A40 | 40 | M | Stomach | Adenocarcinoma | 3 | IIIA | Malignant | 2.5 |
| | A41 | 70 | M | Lymph node | Metastatic adenocarcinoma of No.A1 | 3 | | Metastasis | 1 |
| | A42 | 71 | M | Lymph node | Metastatic adenocarcinoma of No.A2 | 3 | | Metastasis | 2 |
| | A43 | 61 | M | Lymph node | Metastatic adenocarcinoma of No.A3 | 2 | | Metastasis | 0 |
| | A44 | 60 | M | Lymph node | Metastatic adenocarcinoma of No.A4 | 2 | | Metastasis | 1.75 |
| | A45 | 54 | M | Lymph node | Metastatic adenocarcinoma of No.A5 | 3 | | Metastasis | 0.25 |
| | A46 | 72 | M | Lymph node | Metastatic adenocarcinoma of No.A6 | 2 | | Metastasis | 0.1 |
| | A47 | 77 | M | Lymph node | Metastatic adenocarcinoma of No.A7 | 2 | | Metastasis | 1.5 |
| | A48 | 49 | F | Lymph node | Metastatic adenocarcinoma of No.A8 | 3 | | Metastasis | 0.25 |
| | A49 | 63 | M | Lymph node | Metastatic adenocarcinoma of No.A9 | 3 | | Metastasis | 1 |
| | A50 | 66 | M | Lymph node | Metastatic adenocarcinoma of No.A10 | 3 | | Metastasis | 1 |
| | A51 | 53 | M | Lymph node | Metastatic adenocarcinoma of No.A11 | 3 | | Metastasis | 1 |
| | A52 | 68 | F | Lymph node | Metastatic adenocarcinoma of No.A12 | 2 | | Metastasis | 1 |
| | A53 | 64 | F | Lymph node | Metastatic adenocarcinoma of No.A13 | 3 | | Metastasis | 0.25 |
| | A54 | 56 | M | Lymph node | Metastatic adenocarcinoma of No.A14 | 3 | | Metastasis | 1 |
| | A55 | 57 | M | Lymph node | Metastatic adenocarcinoma of No.A15 | 3 | | Metastasis | 0.25 |
| | A56 | 47 | F | Lymph node | Metastatic adenocarcinoma of No.A16 | 2 | | Metastasis | 0 |
| | A57 | 68 | M | Lymph node | Metastatic adenocarcinoma of No.A17 | 2 | | Metastasis | 1 |
| | A58 | 56 | M | Lymph node | Metastatic adenocarcinoma of No.A18 | 3 | | Metastasis | 0 |
| | A58 | 62 | M | Lymph node | Metastatic adenocarcinoma of No.A19 | 3 | | Metastasis | 1 |
| | A59 | 68 | F | Lymph node | Metastatic adenocarcinoma of No.A20 | 3 | | Metastasis | 2 |
| | A60 | 36 | M | Lymph node | Metastatic adenocarcinoma of No.A21 | 3 | | Metastasis | 1 |
| | A61 | 53 | M | Lymph node | Metastatic adenocarcinoma of No.A22 | 3 | | Metastasis | 1.75 |
| | A62 | 54 | M | Lymph node | Metastatic adenocarcinoma of No.A23 | 3 | | Metastasis | 0 |
| | A63 | 34 | M | Lymph node | Metastatic adenocarcinoma of No.A24 | 3 | | Metastasis | 0.1 |
| | A64 | 45 | M | Lymph node | Metastatic adenocarcinoma of No.A25 | 3 | | Metastasis | 0.75 |
| | A65 | 49 | M | Lymph node | Metastatic adenocarcinoma of No.A26 | 2--3 | | Metastasis | 0.25 |
| | A66 | 57 | M | Lymph node | Metastatic adenocarcinoma of No.A27 | 3 | | Metastasis | 1 |
| | A67 | 63 | M | Lymph node | Metastatic adenocarcinoma of No.A28 | 3 | | Metastasis | 1 |
| | A68 | 59 | M | Lymph node | Metastatic adenocarcinoma of No.A29 | 3 | | Metastasis | 0.25 |
| | A69 | 60 | M | Lymph node | Metastatic adenocarcinoma of No.A30 | 3 | | Metastasis | 0.5 |
| | A70 | 47 | F | Lymph node | Metastatic adenocarcinoma of No.A31 | 3 | | Metastasis | 0 |
| | A71 | 50 | M | Lymph node | Metastatic adenocarcinoma of No.A32 | 3 | | Metastasis | 0.1 |
| | A72 | 63 | M | Lymph node | Metastatic adenocarcinoma of No.A33 | 2 | | Metastasis | 0.25 |
| | A73 | 37 | F | Lymph node | Metastatic adenocarcinoma of No.A34 | 3 | | Metastasis | 0 |
| | A74 | 35 | M | Lymph node | Metastatic adenocarcinoma of No.A35 | 2--3 | | Metastasis | 0 |
| | A75 | 68 | M | Lymph node | Metastatic mucinous adenocarcinoma of No.A36 | 3 | | Metastasis | 0 |
| | A76 | 58 | M | Lymph node | Metastatic adenocarcinoma of No.A37 | 3 | | Metastasis | 0 |
| | A77 | 24 | F | Lymph node | Metastatic adenocarcinoma of No.A38 | 2--3 | | Metastasis | 0 |
| | A78 | 64 | M | Lymph node | Metastatic adenocarcinoma of No.A39 | 3 | | Metastasis | 0 |
| | A79 | 40 | M | Lymph node | Metastatic adenocarcinoma of No.A40 | 3 | | Metastasis | 1 |
| **ST8013B** | | | | | | | | | |
| | B1 | 65 | F | Greater omentum | Metastatic adenocarcinoma from stomach | 3 | | Metastasis | 0.75 |
| | B2 | 65 | F | Greater omentum | Metastatic adenocarcinoma from stomach | 2 | | Metastasis | 0.5 |

| Microarray | No. | Age | Sex | Organ/Anatomic Site | Pathology diagnosis | Grade | Stage | Type | Score |
|---|---|---|---|---|---|---|---|---|---|
| | B3 | 50 | M | Greater omentum | Metastatic adenocarcinoma from stomach | 2 | | Metastasis | 0 |
| | B4 | 50 | M | Greater omentum | Metastatic adenocarcinoma from stomach | 2 | | Metastasis | 1 |
| | B5 | 49 | F | Greater omentum | Metastatic mucinous adenocarcinoma from stomach | 3 | | Metastasis | N/A |
| | B6 | 49 | F | Greater omentum | Metastatic mucinous adenocarcinoma from stomach | 3 | | Metastasis | N/A |
| | B7 | 46 | M | Greater omentum | Metastatic adenocarcinoma from stomach | 3 | | Metastasis | 0 |
| | B8 | 46 | M | Greater omentum | Metastatic adenocarcinoma from stomach | 3 | | Metastasis | 0 |
| | B9 | 52 | F | Liver | Metastatic adenocarcinoma from stomach | 2 | | Metastasis | 2.5 |
| | B10 | 52 | F | Liver | Metastatic adenocarcinoma from stomach | 2 | | Metastasis | 2.25 |
| | B11 | 72 | F | Liver | Metastatic adenocarcinoma from stomach | 3 | | Metastasis | 1.5 |
| | B12 | 72 | F | Liver | Metastatic adenocarcinoma from stomach | 3 | | Metastasis | 2.75 |
| | B13 | 45 | M | Liver | Metastatic adenocarcinoma from stomach | 2 | | Metastasis | 2 |
| | B14 | 45 | M | Liver | Metastatic adenocarcinoma from stomach | 2 | | Metastasis | 1 |
| | B15 | 62 | F | Liver | Metastatic adenocarcinoma from stomach | 3 | | Metastasis | 1 |
| | B16 | 62 | F | Liver | Metastatic adenocarcinoma from stomach | 3 | | Metastasis | 0.5 |
| | B17 | 41 | F | Ovary | Metastatic adenocarcinoma from stomach | 3 | | Metastasis | 1 |
| | B18 | 41 | F | Ovary | Metastatic adenocarcinoma from stomach | 3 | | Metastasis | 1 |
| | B19 | 42 | F | Ovary | Metastatic adenocarcinoma from stomach | 3 | | Metastasis | 2.75 |
| | B20 | 42 | F | Ovary | Metastatic adenocarcinoma from stomach | 3 | | Metastasis | 3 |
| | B21 | 62 | M | Ovary | Metastatic adenocarcinoma from stomach | 3 | | Metastasis | 1 |
| | B22 | 62 | M | Ovary | Metastatic adenocarcinoma from stomach | 3 | | Metastasis | 1 |
| | B23 | 75 | M | Lymph node | Metastatic adenocarcinoma | 2 | | Metastasis | 3 |
| | B24 | 75 | M | Lymph node | Metastatic adenocarcinoma | 2 | | Metastasis | 2 |
| | B25 | 50 | M | Lymph node | Metastatic adenocarcinoma | 2 | | Metastasis | 1.5 |
| | B26 | 50 | M | Lymph node | Metastatic adenocarcinoma | 2 | | Metastasis | 0 |
| | B27 | 45 | M | Lymph node | Metastatic adenocarcinoma | 3 | | Metastasis | 0 |
| | B28 | 45 | M | Lymph node | Metastatic adenocarcinoma | 3 | | Metastasis | 1.5 |
| | B29 | 57 | M | Lymph node | Metastatic adenocarcinoma | 3 | | Metastasis | 1 |
| | B30 | 57 | M | Lymph node | Metastatic adenocarcinoma | 3 | | Metastasis | 1.5 |
| | B31 | 63 | M | Lymph node | Metastatic adenocarcinoma | 2 | | Metastasis | 2.5 |
| | B32 | 63 | M | Lymph node | Metastatic adenocarcinoma | 2 | | Metastasis | 0.25 |
| | B33 | 51 | F | Lymph node | Metastatic neuroendocrine carcinoma | 3 | | Metastasis | 0 |
| | B34 | 51 | F | Lymph node | Metastatic neuroendocrine carcinoma | 3 | | Metastasis | 1 |
| | B35 | 62 | M | Lymph node | Metastatic adenocarcinoma | 2 | | Metastasis | 0.5 |
| | B36 | 62 | M | Lymph node | Metastatic adenocarcinoma | 2 | | Metastasis | 2 |
| | B37 | 53 | M | Lymph node | Metastatic adenocarcinoma (sparse) | 3 | | Metastasis | 0.5 |
| | B38 | 53 | M | Lymph node | Metastatic adenocarcinoma (lymph node) | * | | Metastasis | 1.5 |
| | B39 | 58 | M | Lymph node | Metastatic adenocarcinoma | 2 | | Metastasis | 2 |
| | B40 | 58 | M | Lymph node | Metastatic adenocarcinoma | 2 | | Metastasis | 1 |
| | B41 | 61 | M | Lymph node | Metastatic adenocarcinoma | 2 | | Metastasis | 1 |
| | B42 | 61 | M | Lymph node | Metastatic adenocarcinoma | 2 | | Metastasis | 1 |
| | B43 | 67 | M | Lymph node | Metastatic adenocarcinoma | 2 | | Metastasis | 1 |
| | B44 | 67 | M | Lymph node | Metastatic adenocarcinoma | 2 | | Metastasis | 1 |
| | B45 | 66 | M | Lymph node | Metastatic adenocarcinoma | 3 | | Metastasis | 1 |
| | B46 | 66 | M | Lymph node | Metastatic adenocarcinoma | 3 | | Metastasis | 1 |
| | B47 | 37 | F | Lymph node | Metastatic adenocarcinoma | 3 | | Metastasis | 1.75 |
| | B48 | 37 | F | Lymph node | Metastatic adenocarcinoma | 3 | | Metastasis | 2 |
| | B49 | 74 | M | Lymph node | Metastatic adenocarcinoma | 3 | | Metastasis | 1.5 |
| | B50 | 74 | M | Lymph node | Metastatic adenocarcinoma | 3 | | Metastasis | 0 |
| | B51 | 57 | M | Lymph node | Metastatic adenocarcinoma | 3 | | Metastasis | 0 |
| | B52 | 57 | M | Lymph node | Metastatic adenocarcinoma | 3 | | Metastasis | 0.5 |
| | B53 | 70 | M | Lymph node | Metastatic adenocarcinoma | 3 | | Metastasis | 0.5 |
| | B54 | 70 | M | Lymph node | Metastatic adenocarcinoma | 3 | | Metastasis | 0.5 |
| | B55 | 45 | M | Lymph node | Metastatic adenocarcinoma | 3 | | Metastasis | 0.25 |
| | B56 | 45 | M | Lymph node | Metastatic adenocarcinoma | 3 | | Metastasis | 0.5 |
| | B57 | 73 | M | Lymph node | Metastatic adenocarcinoma | 3 | | Metastasis | 0.25 |
| | B58 | 73 | M | Lymph node | Metastatic adenocarcinoma | 3 | | Metastasis | 0 |
| | B59 | 48 | M | Lymph node | Metastatic adenocarcinoma | 3 | | Metastasis | 2.95 |
| | B60 | 48 | M | Lymph node | Metastatic adenocarcinoma (sparse) | 3 | | Metastasis | 3 |
| | B61 | 53 | M | Lymph node | Metastatic adenocarcinoma | 3 | | Metastasis | 2 |
| | B62 | 53 | M | Lymph node | Metastatic adenocarcinoma | 3 | | Metastasis | 1.5 |
| | B63 | 48 | M | Lymph node | Metastatic adenocarcinoma (sparse) | 3 | | Metastasis | 0.25 |
| | B64 | 48 | M | Lymph node | Metastatic adenocarcinoma with necrosis | 3 | | Metastasis | 0 |
| | B65 | 67 | M | Lymph node | Metastatic adenocarcinoma with necrosis | 3 | | Metastasis | 1 |
| | B66 | 67 | M | Lymph node | Metastatic adenocarcinoma with necrosis | 3 | | Metastasis | 1 |
| | B67 | 61 | M | Lymph node | Metastatic adenocarcinoma | 3 | | Metastasis | 2 |
| | B68 | 61 | M | Lymph node | Metastatic adenocarcinoma | 3 | | Metastasis | 1 |
| | B69 | 34 | F | Lymph node | Metastatic adenocarcinoma | 3 | | Metastasis | 0.25 |
| | B70 | 34 | F | Lymph node | Metastatic adenocarcinoma | 3 | | Metastasis | 0.25 |
| | B71 | 52 | M | Lymph node | Metastatic adenocarcinoma | 3 | | Metastasis | 0.5 |
| | B72 | 52 | M | Lymph node | Metastatic adenocarcinoma | 3 | | Metastasis | 1.5 |
| | B73 | 53 | M | Greater omentum | Metastatic adenocarcinoma | 2 | | Metastasis | 1 |
| | B74 | 53 | M | Greater omentum | Metastatic adenocarcinoma | 2 | | Metastasis | 0.5 |
| | B75 | 55 | M | Lymph node | Metastatic adenocarcinoma | 3 | | Metastasis | 1.5 |
| | B76 | 55 | M | Lymph node | Metastatic adenocarcinoma | 3 | | Metastasis | 1 |
| | B77 | 50 | M | Lymph node | Metastatic adenocarcinoma | 3 | | Metastasis | 3 |
| | B78 | 50 | M | Lymph node | Metastatic adenocarcinoma | 3 | | Metastasis | 2.75 |
| | B79 | 58 | M | Lymph node | Metastatic adenocarcinoma | 3 | | Metastasis | 2.5 |
| **ST1451A** | | | | | | | | | |
| | C1 | 30 | F | Stomach | Adenocarcinoma | 1 | IIIB | Malignant | 1 |
| | C2 | 48 | F | Stomach | Adenocarcinoma | 1 | IIIA | Malignant | 1 |
| | C3 | 63 | M | Stomach | Adenocarcinoma (smooth muscle) | * | IB | Malignant | 0 |
| | C4 | 64 | M | Stomach | Adenocarcinoma | 2 | IIA | Malignant | 1 |
| | C5 | 59 | M | Stomach | Adenocarcinoma (stomach tissue) | * | IIA | Malignant | 0.25 |
| | C6 | 52 | M | Stomach | Adenocarcinoma | 1 | IIB | Malignant | 2 |

| Microarray | No. | Age | Sex | Organ/Anatomic Site | Pathology diagnosis | Grade | Stage | Type | Score |
|---|---|---|---|---|---|---|---|---|---|
| | C7 | 61 | M | Stomach | Adenocarcinoma | 2 | IIA | Malignant | 1.5 |
| | C8 | 59 | M | Stomach | Adenocarcinoma | 2 | IIA | Malignant | N/A |
| | C9 | 58 | M | Stomach | Adenocarcinoma | 2 | IIIA | Malignant | 0.5 |
| | C10 | 37 | F | Stomach | Adenocarcinoma (fibrous tissue) | * | IIIB | Malignant | 0 |
| | C11 | 52 | M | Stomach | Adenocarcinoma | 2 | IIA | Malignant | 0.5 |
| | C12 | 32 | M | Stomach | Adenocarcinoma | 2 | IIIA | Malignant | 0 |
| | C13 | 79 | M | Stomach | Adenocarcinoma (sparse) | 2 | IIA | Malignant | 0 |
| | C14 | 56 | F | Stomach | Adenocarcinoma | 2 | IIA | Malignant | 1 |
| | C15 | 45 | M | Stomach | Adenocarcinoma | 2 | IIB | Malignant | 2 |
| | C16 | 56 | F | Stomach | Adenocarcinoma | 1 | IIIA | Malignant | 2.25 |
| | C17 | 67 | M | Stomach | Adenocarcinoma | 3 | IIIA | Malignant | 0.25 |
| | C18 | 62 | M | Stomach | Adenocarcinoma | 2 | IIIA | Malignant | 2 |
| | C19 | 67 | M | Stomach | Adenocarcinoma | 2 | IIA | Malignant | 1 |
| | C20 | 63 | M | Stomach | Adenocarcinoma (stomach tissue) | * | IIIB | Malignant | 0.25 |
| | C21 | 65 | M | Stomach | Adenocarcinoma | 2 | IIA | Malignant | 1 |
| | C22 | 62 | M | Stomach | Adenocarcinoma | 2 | IIIA | Malignant | 1.5 |
| | C23 | 51 | M | Stomach | Adenocarcinoma | 2--3 | IIIC | Malignant | 2 |
| | C24 | 59 | M | Stomach | Adenocarcinoma invades nerve | 3 | IIA | Malignant | 0.5 |
| | C25 | 57 | M | Stomach | Adenocarcinoma | 2 | IIB | Malignant | 1 |
| | C26 | 42 | M | Stomach | Adenocarcinoma | 3 | IIIC | Malignant | 1.75 |
| | C27 | 42 | M | Stomach | Adenocarcinoma | 3 | IIA | Malignant | 0.25 |
| | C28 | 41 | M | Stomach | Adenocarcinoma (sparse) | 3 | IIIC | Malignant | N/A |
| | C29 | 50 | M | Stomach | Adenocarcinoma | 3 | IIA | Malignant | 0.25 |
| | C30 | 43 | M | Stomach | Adenocarcinoma | 3 | IIA | Malignant | 0.25 |
| | C31 | 47 | M | Stomach | Adenocarcinoma | 3 | IIA | Malignant | 0 |
| | C32 | 44 | F | Stomach | Adenocarcinoma | 3 | IIA | Malignant | 0.25 |
| | C33 | 59 | M | Stomach | Adenocarcinoma (smooth muscle) | * | IIA | Malignant | 0.5 |
| | C34 | 69 | M | Stomach | Adenocarcinoma (smooth muscle) | * | IIA | Malignant | 1 |
| | C35 | 38 | M | Stomach | Adenocarcinoma | 3 | IIA | Malignant | 1.5 |
| | C36 | 67 | M | Stomach | Adenocarcinoma (smooth muscle) | * | IIB | Malignant | N/A |
| | C37 | 72 | M | Stomach | Adenocarcinoma | 3 | IIA | Malignant | 2.5 |
| | C38 | 57 | M | Stomach | Adenocarcinoma | 3 | IIB | Malignant | 2 |
| | C39 | 55 | M | Stomach | Adenocarcinoma | 3 | IIA | Malignant | 0 |
| | C40 | 52 | M | Stomach | Adenocarcinoma | 2 | IIB | Malignant | 1.25 |
| | C41 | 63 | F | Stomach | Adenocarcinoma | 3 | IIB | Malignant | 2.75 |
| | C42 | 60 | M | Stomach | Adenocarcinoma (fibrous tissue and blood vessel) | * | IIIB | Malignant | 1.25 |
| | C43 | 63 | M | Stomach | Mucinous adenocarcinoma | 3 | IIIA | Malignant | N/A |
| | C44 | 47 | M | Stomach | Adenocarcinoma (fibrous tissue and blood vessel) | * | IIIB | Malignant | N/A |
| | C45 | 62 | M | Stomach | Adenocarcinoma | 3 | IIIA | Malignant | 1.25 |
| | C46 | 58 | M | Stomach | Adenocarcinoma | 3 | IIA | Malignant | 0.75 |
| | C47 | 73 | M | Stomach | Adenocarcinoma (fibrous tissue and blood vessel) | * | IIA | Malignant | N/A |
| | C48 | 44 | F | Stomach | Adenocarcinoma (smooth muscle and fibrou tissue) | * | IB | Malignant | N/A |
| | C49 | 59 | M | Stomach | Adenocarcinoma | 3 | IIB | Malignant | 2 |
| | C50 | 62 | M | Stomach | Adenocarcinoma | 3 | IIA | Malignant | 1 |
| | C51 | 63 | M | Stomach | Adenocarcinoma | 3 | IIIA | Malignant | 1.25 |
| | C52 | 52 | M | Stomach | Adenocarcinoma | 3 | IIIB | Malignant | 0.25 |
| | C53 | 73 | M | Stomach | Adenocarcinoma | 3 | IIB | Malignant | 2.5 |
| | C54 | 53 | M | Stomach | Adenocarcinoma | 3 | IIB | Malignant | 0 |
| | C55 | 53 | F | Stomach | Adenocarcinoma | 3 | IB | Malignant | 2.75 |
| | C56 | 63 | F | Stomach | Adenocarcinoma (smooth muscle) | * | IIB | Malignant | 0.25 |
| | C57 | 54 | M | Stomach | Adenocarcinoma | 3 | IIIB | Malignant | 1.25 |
| | C58 | 61 | M | Stomach | Adenocarcinoma (smooth muscle) | * | IIIC | Malignant | 2.5 |
| | C59 | 68 | F | Stomach | Adenocarcinoma (smooth muscle) | * | IIIA | Malignant | 3 |
| | C60 | 47 | M | Stomach | Adenocarcinoma | 2 | IIIC | Malignant | 0 |
| | C61 | 44 | M | Stomach | Adenocarcinoma (smooth muscle) | * | IIB | Malignant | N/A |
| | C62 | 68 | M | Stomach | Mucinous adenocarcinoma | 3 | IIIA | Malignant | N/A |
| | C63 | 72 | M | Stomach | Mucinous adenocarcinoma | 3 | IIIA | Malignant | 1.75 |
| | C64 | 61 | F | Stomach | Mucinous adenocarcinoma | 3 | IIIA | Malignant | 2 |
| | C65 | 42 | M | Stomach | Mucinous adenocarcinoma | 3 | IIIA | Malignant | 2.5 |
| | C66 | 71 | M | Stomach | Signet-ring cell carcinoma | - | IIA | Malignant | 0.25 |
| | C67 | 61 | F | Stomach | Signet-ring cell carcinoma | - | IIA | Malignant | 0 |
| | C68 | 68 | M | Stomach | Signet-ring cell carcinoma | - | IIA | Malignant | 0.5 |
| | C69 | 61 | M | Stomach | Neuroendocrine carcinoma | - | IIB | Malignant | 1 |
| | C70 | 66 | M | Stomach | Neuroendocrine carcinoma | - | IIA | Malignant | 2.75 |
| | C71 | 30 | F | Stomach | Adjacent normal stomach tissue of No.C1 (fibrous tissue and blood vessel) | | | NAT | 0.5 |
| | C72 | 48 | F | Stomach | Adjacent normal stomach tissue of No.C2 | | | NAT | 2.5 |
| | C73 | 63 | M | Stomach | Adjacent normal stomach tissue of No.C3 | | | NAT | 1 |
| | C74 | 64 | M | Stomach | Cancer adjacent stomach tissue of No.C4 | | | AT | 3 |
| | C75 | 59 | M | Stomach | Adjacent normal stomach tissue of No.C5 (fibrous tissue and blood vessel) | | | NAT | 1.5 |
| | C76 | 52 | M | Stomach | Adjacent normal stomach tissue of No.C6 | | | NAT | 2.25 |
| | C77 | 61 | M | Stomach | Adjacent normal stomach tissue of No.C7 | | | NAT | 1.75 |
| | C78 | 59 | M | Stomach | Adjacent normal stomach tissue of No.C8 | | | NAT | 2.75 |
| | C79 | 58 | M | Stomach | Adjacent normal stomach tissue of No.C9 (fibrous and fatty tissue) | | | NAT | 2.5 |
| | C80 | 37 | F | Stomach | Adjacent normal stomach tissue of No.C10 | | | NAT | 2.5 |
| | C81 | 52 | M | Stomach | Adjacent normal stomach tissue of No.C11 | | | NAT | 2.5 |
| | C82 | 32 | M | Stomach | Adjacent normal stomach tissue of No.C12 | | | NAT | 1 |
| | C83 | 79 | M | Stomach | Cancer adjacent stomach tissue of No.C13 | | | AT | N/A |
| | C84 | 56 | F | Stomach | Adjacent normal stomach tissue of No.C14 | | | NAT | 2.5 |
| | C85 | 45 | M | Stomach | Adjacent normal stomach tissue of No.C15 (fibrous tissue and blood vessel) | | | NAT | 2 |
| | C86 | 56 | F | Stomach | Cancer adjacent stomach tissue of No.C16 | | | AT | 2 |
| | C87 | 67 | M | Stomach | Adjacent normal stomach tissue of No.C17 (fibrous tissue and blood vessel) | | | NAT | 2.25 |
| | C88 | 62 | M | Stomach | Adjacent normal stomach tissue of No.C18 | | | NAT | 1 |
| | C89 | 67 | M | Stomach | Adjacent normal stomach tissue of No.C19 (fibrous tissue and blood vessel) | | | NAT | 1.5 |
| | C90 | 63 | M | Stomach | Adjacent normal stomach tissue of No.C20 | | | NAT | 0.5 |

| Microarray | No. | Age | Sex | Organ/Anatomic Site | Pathology diagnosis | Grade | Stage | Type | Score |
|---|---|---|---|---|---|---|---|---|---|
| | C91 | 65 | M | Stomach | Adjacent normal stomach tissue of No.C21 | | | NAT | 2.5 |
| | C92 | 62 | M | Stomach | Adjacent normal stomach tissue of No.C22 | | | NAT | 2 |
| | C93 | 51 | M | Stomach | Adjacent normal stomach tissue of No.C23 | | | NAT | 2 |
| | C94 | 59 | M | Stomach | Adjacent normal stomach tissue of No.C24 | | | NAT | 2 |
| | C95 | 57 | M | Stomach | Adjacent normal stomach tissue of No.C25 | | | NAT | 2.25 |
| | C96 | 42 | M | Stomach | Adjacent normal stomach tissue of No.C26 | | | NAT | 1 |
| | C97 | 42 | M | Stomach | Adjacent normal stomach tissue of No.C27 | | | NAT | N/A |
| | C98 | 41 | M | Stomach | Adjacent normal stomach tissue of No.C28 | | | NAT | N/A |
| | C99 | 50 | M | Stomach | Adjacent normal stomach tissue of No.C29 | | | NAT | 3 |
| | C100 | 43 | M | Stomach | Adjacent normal stomach tissue of No.C30 | | | NAT | 3 |
| | C101 | 47 | M | Stomach | Cancer adjacent stomach tissue of No.C31 | | | AT | N/A |
| | C102 | 44 | F | Stomach | Adjacent normal stomach tissue of No.C32 | | | NAT | 2.75 |
| | C103 | 59 | M | Stomach | Adjacent normal stomach tissue of No.C33 | | | NAT | 1.5 |
| | C104 | 69 | M | Stomach | Adjacent normal stomach tissue of No.C34 | | | NAT | 1 |
| | C105 | 38 | M | Stomach | Adjacent normal stomach tissue of No.C35 | | | NAT | 1 |
| | C106 | 67 | M | Stomach | Adjacent normal stomach tissue of No.C36 | | | NAT | 1.5 |
| | C107 | 72 | M | Stomach | Cancer adjacent stomach tissue of No.C37 | | | AT | 0.5 |
| | C108 | 57 | M | Stomach | Cancer adjacent stomach tissue of No.C38 | | | AT | 3 |
| | C109 | 55 | M | Stomach | Adjacent normal stomach tissue of No.C39 | | | NAT | 3 |
| | C110 | 52 | M | Stomach | Adjacent normal stomach tissue of No.C40 | | | NAT | 3 |
| | C111 | 63 | F | Stomach | Adjacent normal stomach tissue of No.C41 | | | NAT | 2.5 |
| | C112 | 60 | M | Stomach | Adjacent normal stomach tissue of No.C42 | | | NAT | 3 |
| | C113 | 63 | M | Stomach | Adjacent normal stomach tissue of No.C43 | | | NAT | 3 |
| | C114 | 47 | M | Stomach | Adjacent normal stomach tissue of No.C44 | | | NAT | 2 |
| | C115 | 62 | M | Stomach | Adjacent normal stomach tissue of No.C45 | | | NAT | 3 |
| | C116 | 58 | M | Stomach | Adjacent normal stomach tissue of No.C46 | | | NAT | 3 |
| | C117 | 73 | M | Stomach | Adjacent normal stomach tissue of No.C47 | | | NAT | 2.75 |
| | C118 | 44 | F | Stomach | Cancer adjacent stomach tissue of No.C48 | | | AT | N/A |
| | C119 | 59 | M | Stomach | Adjacent normal stomach tissue of No.C49 | | | NAT | 0.5 |
| | C120 | 62 | M | Stomach | Cancer adjacent stomach tissue of No.C50 | | | AT | N/A |
| | C121 | 63 | M | Stomach | Adjacent normal stomach tissue of No.C51 | | | NAT | 2 |
| | C122 | 52 | M | Stomach | Cancer adjacent stomach tissue of No.C52 | | | AT | N/A |
| | C123 | 73 | M | Stomach | Adjacent normal stomach tissue of No.C53 | | | NAT | 2.25 |
| | C124 | 53 | M | Stomach | Adjacent normal stomach tissue of No.C54 | | | NAT | 2 |
| | C125 | 53 | F | Stomach | Adjacent normal stomach tissue of No.C55 | | | NAT | 3 |
| | C126 | 63 | F | Stomach | Cancer adjacent stomach tissue of No.C56 | | | AT | N/A |
| | C127 | 54 | M | Stomach | Adjacent normal stomach tissue of No.C57 | | | NAT | 2 |
| | C128 | 61 | M | Stomach | Adjacent normal stomach tissue of No.C58 | | | NAT | 2 |
| | C129 | 68 | F | Stomach | Adjacent normal stomach tissue of No.C59 | | | NAT | N/A |
| | C130 | 47 | M | Stomach | Adjacent normal stomach tissue of No.C60 | | | NAT | 0 |
| | C131 | 44 | M | Stomach | Cancer adjacent stomach tissue of No.C61 | | | AT | 1 |
| | C132 | 68 | M | Stomach | Adjacent normal stomach tissue of No.C62 | | | NAT | 1.75 |
| | C133 | 72 | M | Stomach | Adjacent normal stomach tissue of No.C63 | | | NAT | 2.25 |
| | C134 | 61 | F | Stomach | Adjacent normal stomach tissue of No.C64 | | | NAT | 1.25 |
| | C135 | 42 | M | Stomach | Adjacent normal stomach tissue of No.C65 | | | NAT | 2.5 |
| | C136 | 71 | M | Stomach | Adjacent normal stomach tissue of No.C66 | | | NAT | 0.25 |
| | C137 | 61 | F | Stomach | Adjacent normal stomach tissue of No.C67 | | | NAT | 2.5 |
| | C138 | 68 | M | Stomach | Adjacent normal stomach tissue of No.C68 | | | NAT | 3 |
| | C139 | 61 | M | Stomach | Adjacent normal stomach tissue of No.C69 | | | NAT | 3 |
| | C140 | 66 | M | Stomach | Adjacent normal stomach tissue of No.C70 | | | NAT | 2.5 |
| | C141 | 58 | M | Lymph node | Metastatic adenocarcinoma from stomach of No.C9 | 3 | | Metastasis | 2.5 |
| | C142 | 44 | M | Lymph node | Metastatic adenocarcinoma from stomach of No.C61 (lymph node) | * | | Metastasis | 3 |
| | C143 | 75 | M | Lymph node | Metastatic adenocarcinoma from stomach | 3 | | Metastasis | 0 |
| | C144 | 61 | M | Lymph node | Metastatic neuroendocrine carcinoma form stomach No.C69 | - | | Metastasis | 0 |
| | C145 | 57 | M | Lymph node | Metastatic adenocarcinoma from stomach of No.C25 | 3 | | Metastasis | 1 |
| **T011B** | | | | | | | | | |
| | D1 | 71 | F | stomach | Adenocarcinoma | 1 | IB | Malignant | 2.5 |
| | D2 | 71 | F | stomach | Adenocarcinoma | 1 | IB | Malignant | 3 |
| | D3 | 60 | M | stomach | Mucinous adenocarcinoma with necrosis | 3 | IIA | Malignant | N/A |
| | D4 | 60 | M | stomach | Mucinous adenocarcinoma | 3 | IIA | Malignant | N/A |
| | D5 | 71 | F | stomach | Adenocarcinoma | 1 | IB | Malignant | 2 |
| | D6 | 71 | F | stomach | Adenocarcinoma | 1 | IB | Malignant | 2.5 |
| | D7 | 60 | M | stomach | Mucinous adenocarcinoma | 3 | IIA | Malignant | 0 |
| | D8 | 60 | M | stomach | Mucinous adenocarcinoma | 3 | IIA | Malignant | N/A |
| | D9 | 90 | M | stomach | Signet ring cell carcinoma | - | IIB | Malignant | 2 |
| | D10 | 90 | M | stomach | Signet ring cell carcinoma | - | IIB | Malignant | 1.5 |
| | D11 | 42 | F | stomach | Squamous cell carcinoma | 3 | IIA | Malignant | 1.5 |
| | D12 | 42 | F | stomach | Signet ring cell carcinoma | - | IIB | Malignant | 1 |
| | D13 | 90 | M | stomach | Signet ring cell carcinoma | - | IIB | Malignant | 0.5 |
| | D14 | 90 | M | stomach | Squamous cell carcinoma | 3 | IIA | Malignant | 0.5 |
| | D15 | 42 | F | Stomach | Squamous cell carcinoma | 3 | IIA | Malignant | N/A |
| | D16 | 42 | F | stomach | Stomach tissue | | | Normal | 3 |
| | D17 | 30 | M | stomach | Stomach tissue | | | Normal | 2.5 |
| | D18 | 30 | M | stomach | Stomach tissue | | | Normal | 2.5 |
| | D19 | 38 | M | stomach | Stomach tissue | | | Normal | 3 |
| | D20 | 38 | M | stomach | Stomach tissue | | | Normal | 2 |
| | D21 | 30 | M | stomach | Stomach tissue | | | Normal | 2 |
| | D22 | 30 | M | stomach | Stomach tissue | | | Normal | 1.5 |
| | D23 | 38 | M | stomach | Stomach tissue | | | Normal | 2.5 |

### Figure S1
